# FFselect: An improved linear mixed model for genome-wide association study in populations featuring shared environments confounded by relatedness

**DOI:** 10.1101/2020.01.01.892455

**Authors:** Nichol Schultz, Kent Weigel

## Abstract

Linear mixed models are effective tools to identify genetic loci contributing to phenotypic variation while handling confounding due to population structure and cryptic relatedness. Recent improvements of the linear mixed model for genome-wide association analysis have been directed at more accurately modeling loci of large effect. We describe FFselect (https://github.com/NicholSchultz/FFselect), a novel method that both builds upon recent advances and further extends the linear mixed model for genome-wide association analysis to allow modeling of shared environmental effects. FFselect improves power, controls false discovery rate, and simultaneously corrects for environmental confounding to improve the utility of GWAS.

## Introduction

Genome-wide association studies (GWAS) have become a common approach to identify genetic loci contributing to phenotypic variation in humans and agriculturally important traits in plants and animals. Single marker tests are commonly performed to test for associations between single nucleotide polymorphisms (SNP) and a trait of interest. However, significant associations may occur for reasons other than true linkage between a causal variant and a test marker. Population structure and cryptic relatedness cause genome-wide linkage disequilibrium between loci that are physically unlinked, resulting in false positive associations (Rosenberg & Nordborg, 2006).

Linear mixed models effectively handle confounding due to the genetic background of causal variants in the presence of population structure by fitting the total genetic effect of individuals as a random polygenic term having a variance and covariance structure defined by a genomic relationship matrix (GRM) so that correlation in phenotype reflects relatedness (Yu et al., 2006). The GRM is derived from the entire set of genetic markers and an infinitesimal model is assumed, i.e. all genetic markers are assumed to have some small effect on the trait. Therefore mixed models adjust well for the confounding effects of a diffuse background due to a large number of loci of small effect, but do not always appropriately account for loci of larger effect.

Recent strategies to improve the power of the linear mixed model for GWAS have been directed at more accurately modeling loci of large effect. Instead of using all of the genetic markers to define relationships among individuals, the FaST-LMM-Select (Lippert et al., 2013; Wang, Tian, Pan, Buckler, & Zhang, 2014) and SUPER (Wang et al., 2014) methods use only select associated genetic markers as “pseudo” quantitative trait nucleotides (QTN). Single marker tests are still performed to test for association with the trait, however pseudo QTN correlated with the test marker are excluded from the selected SNP used to derive the GRM (Lippert et al., 2013; Wang et al., 2014). In the FaST-LMM-Select method, a pseudo QTN is considered correlated if it is within a specified physical distance on either side of the test marker whereas the SUPER method applies a threshold on linkage disequilibrium between the pseudo QTN and the test marker. This strategy of selectively including pseudo QTN to derive the GRM for a specific testing marker has been shown to improve statistical power compared to a GRM determined from all or a random sample of SNP (Lippert et al., 2013; Wang et al., 2014; Widmer et al., 2014).

The strategy described above increases statistical power by more accurately modeling the impact of large effect loci on the random polygenic term. Alternatively, a similar approach has been proposed that fits pseudo QTN as fixed effects in addition to the testing markers in a stepwise Multi-Locus Mixed-Model (MLMM) (Segura et al., 2012). The FarmCPU (Liu, Huang, Fan, Buckler, & Zhang, 2016) method also fits pseudo QTN as fixed covariates after an iterative selection process that involves identifying the pseudo QTN that maximize the model likelihood when used to derive the GRM in the random effect model. With the exception of the MLMM method, the described methods do not retain a GRM derived from the entire set of genetic markers. (Widmer et al., 2014) demonstrated that a GRM derived from a select subset of markers does not always adequately correct for familial structure and proposed using a GRM consisting of a mixture of two component GRMs, one constructed from all SNP and another constructed from SNP that well predict the phenotype which they showed to control type I error and yield more power than the standard LMM.

Moreover, familial structure is often spatially confounded, i.e. individuals who share a common environment are likely to be more genetically similar than individuals from two different environments. Phenotypes collected from individuals that share a common environment are typically pre-adjusted for common environmental effects before testing for phenotypic associations with genetic markers. However, this correction, prior to performing the GWAS, may also result in the simultaneous and unwanted removal of a proportion of the genetic variance.

The objective of the current study was to build upon recent improvements of linear mixed models for genome-wide association analysis. We propose a novel method that improves power, controls false discovery rate, and simultaneously corrects for environmental confounding to improve the utility of GWAS.

## Materials and Methods

### FFselect Method: GWAS model

The FFselect (forward feature selection) method was developed in the framework of a standard linear mixed model approach, which decomposes the observation (**Y**) into fixed effect (**β**), random effect (**u**) and residual (**e**) as follows:

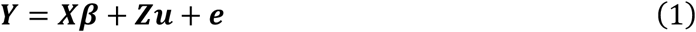

Where **u** is a vector of size **n**(number of individuals) for unknown random effects having a distribution with mean of zero and covariance matrix equal to 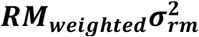, where **RM** is a relationship matrix consisting of a mixture of three component relationship matrices. Relationship Matrix 1 (RM1) is an environmental relationship matrix with element RM1_ij_ (i, j = 1, 2, …, n) equal to 1 for individuals who share the same environment and 0 for individuals from different environments. Relationship Matrix 2 (RM2) is a genomic relationship matrix with element RM2_ij_ (i, j = 1, 2, …, n) calculated from a subset of select genetic markers. Relationship Matrix 3 (RM3) is a genomic relationship matrix with element RM3_ij_ (i, j = 1, 2, …, n) calculated from all genetic markers. RM2 and RM3 are centered genomic relationship matrices calculated by multiplying the centered genotype matrix by its transpose and dividing by the number of markers. RM1, RM2, and RM3 are weighted by their contributions to the unknown variance (**σ_rm_^2^**). **X** and **Z** are the incidence matrices for **β** and **u**, respectively, and random residual effects **e** are normally distributed with zero mean and covariance equal to 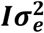, where **I** is the identity matrix and (**σ_e_^2^**) is the unknown residual variance. The random effect **u** is equivalent to the summed environmental and additive genetic effects. For GWAS purposes, it is unnecessary to explicitly estimate each of these effects. Determination of RM1, RM2, and RM3 weights is described further below.

Solving equation (1) involves determining the unknown parameters under which the observations (**Y**) have the maximum log-likelihood using the R package lrgpr (Hoffman, Mezey, & Schadt, 2014) and efficient algorithms of (Lippert et al., 2011) and (Listgarten et al., 2012) The relationship matrix enters the linear mixed model only through its spectral decomposition.

To perform a GWAS, marker effect (**v**) is added to equation (1) one at a time:

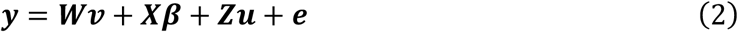

Where **W** is the incidence matrix for **v**. The lrgpr software provides a computational time-saving option of solving equation (2) by using P3D (Zhang et al., 2010) or EMMAX (Kang et al., 2010) which estimates (**σ_rm_^2^**) and (**σ_e_^2^**) once and reuses the estimates (or the ratio **σ_e_^2^**)/**σ_rm_^2^**) for the optimization of **v** and **β** for each genetic test marker.

Proximal contamination is avoided during single marker tests by excluding markers from the select GRM (RM2) that are within a specified base pair distance (e.g. 500 KB was used in both the mice and cattle datasets to achieve desired mapping precision) of the testing marker.

### FFselect Method: Identification of select feature SNP

Identification of select feature SNP to be included in RM2 is performed in a forward stepwise feature selection process. Phenotypes are first adjusted for an estimated shared environment effect determined from a model that includes fixed effect covariates and a random polygenic effect. The adjusted phenotype is next used in a stepwise linear regression model, similar as previously described by (Li, Su, Jiang, & Bao, 2017) and (Segura et al., 2012) A standard LMM GWAS is performed and the most significant SNP is selected for inclusion as a fixed cofactor with each regression step. The polygenic and residual variance components are re-estimated by maximum likelihood at each regression step and the forward selection process continues until the null model polygenic variance estimate is reduced by 95%. In order to prevent overfitting of the data, additional criteria for an earlier forward selection stopping point include selecting the model with the lowest Bayesian Information Criterion (BIC) (Schwarz, 1978). In addition, SNP adjacent (3 SNP upstream and downstream) to the feature selected SNP are also included in the select RM2 to better approximate relationships in the region. In an effort to improve computationally efficiency, only SNP with an initial p-value less than 0.05 proceed through the stepwise selection process.

### FFselect Method: Optimization of RM1, RM2, and RM3 weights

After the feature SNP have been selected for inclusion in RM2, the optimal weights for RM1 (environmental relationship matrix), RM2 (select SNP relationship matrix), and RM3 (all SNP relationship matrix) are determined for use in the final GWAS model. A Brent’s optimization (Brent, 1971) is first performed to maximize the log-likelihood and determine the optimal RM2 and RM3 weights for the phenotype pre-adjusted for the shared environment effect. These weights are then used for a combined RM2 and RM3 matrix when estimating the RM1 weight by Brent’s optimization for the unadjusted phenotype. The results of this two-step approach are very similar to an approach that simultaneously estimates all three weights, however the two-step simplified approach is more computationally efficient.

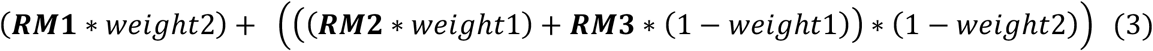

### Real genotype data

The previously published mouse genotype dataset (Neves, Carvalheiro, & Queiroz, 2012) is a heterogeneous stock population owned by the Welcome Trust Centre for Human Genetics. The dataset analyzed in this study was comprised of 1814 mice housed in 523 cages of full-sibs and genotypes at 8850 unique SNP with a minor allele frequency greater than 5%. The dairy cattle genotype dataset includes 3417 Holsteins from 9 United States herds and a single Canadian herd genotyped with 56,483 SNP with minor allele frequency greater than 5%.

### Simulated phenotypes

Real genotype datasets from mice and Holstein dairy cattle were used to simulate genetic effects and generate phenotypes by adding environmental and residual effects. The QTN underlying these simulated phenotypes were randomly sampled from the real genotypes. The QTN effects followed a Gaussian distribution. The phenotype was simulated as: y = additive effect + shared environment effect + residual effect. The additive effect was calculated as: additive effect = QTN matrix * QTN effects. The residual effect followed a Gaussian distribution with mean of 0 and variance equivalent to 0.8*additive genetic variance. The shared environmental effect followed a Gaussian distribution with a mean of 0 and variance equivalent to 0.2*additive genetic variance. Simulations were performed using 30, 100, and 300 QTN with a heritability of 0.5. The QTN were excluded from the genotypic data for association tests. Simulations were replicated 100 times.

### Variance decomposition

Three different linear mixed models were assessed for their ability to accurately estimate the simulated environmental variance (0.1), genetic variance (0.5), and residual variance (0.4). Model 1 included only a random environment effect, Model 2 included only a random genetic effect, and Model 3 included both random environment and random genetic effects.

### Power, type 1 error, and FDR examination

Statistical power, Type I error, and FDR were examined at p-value threshold cutoff values at a strict Bonferroni level for three methods: 1) standard linear mixed model, 2) FarmCPU (Liu et al., 2016), and 3) the method detailed above which we will refer to as “FFselect.” A QTN was considered identified if a positive marker was within 2.5 MB. A distance of 2.5 MB was used to limit the number of false positives generated by the standard linear mixed model due to linkage disequilibrium with large effect causal loci. Power was defined as the proportion of QTN identified. The null distribution of Type I error was derived from non-QTN markers (markers not within 2.5 MB of a QTN). FDR was defined as the proportion of the non-QTN markers among the positive markers.

## Results

Synthetic phenotypes were simulated by adding both genetic effects to real genotype data in addition to shared environment effects for a mice population where full-sibs shared cages and Holstein dairy cattle from nine US herds and one Canadian herd. Additive genetic effects were simulated for a 30, 100, and 300 locus model with environmental, genetic, and residual variance equal to 0.1, 0.5, and 0.4, respectively. Causal loci were removed from the GWAS analysis.

The results for variance decomposition estimates for three different linear mixed models are shown in Table 1. Model 1 includes only a random environment effect, Model 2 includes only a random genetic effect, and Model 3 includes both random environment and random genetic effects. Model 1 overestimates the environmental variance in mice, whereas Model 3 accurately estimates both the environment and genetic variance component in mice and cattle. GWAS models that pre-adjust for shared environmental effects without considering environment may be confounded by genetic relatedness, may be problematic and result in simultaneous removal of a portion of the genetic variance with correction for shared environmental effects. Our proposed forward feature selection method (FFselect) circumvents this issue by including both random environment and genetic effects in the GWAS model.

**Table 1.**
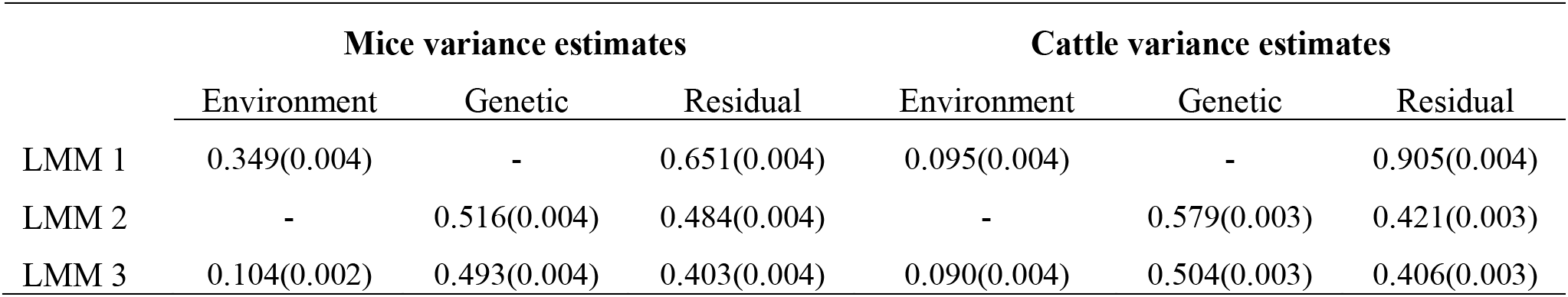
Mean (standard error) variance estimates for 100 simulation replicates of a 100 QTN trait with simulated environmental variance=0.1, genetic variance=0.5, and residual variance=0.4. Linear mixed model (LMM) 1 includes a random environment effect, LMM 2 includes a random genetic effect, and LMM 3 includes both random environment and genetic effects.

We compared our FFselect method with two other linear mixed model mapping methods, a standard method that does not take into account the effects of other major loci or shared environment and a method (FarmCPU) that takes into account the effects of major loci but does not consider loci with small genetic effects or shared environmental effects. The FarmCPU method was selected based on its reported superiority in terms of statistical power and computational efficiency in comparison to other SNP feature selection methods (Liu et al., 2016). Phenotypes were pre-adjusted for environmental effects prior to the standard LMM and FarmCPU analysis. Comparison of statistical power, false discovery rate (FDR) and type I error rate are presented in Table 2 and Figure 1, and can be summarized as follows. First, the methods that take into account the effects of other major loci demonstrate improved power. However, this comes at the cost of a high FDR, unless the polygenic background is also included in the model.

**Table 2.**
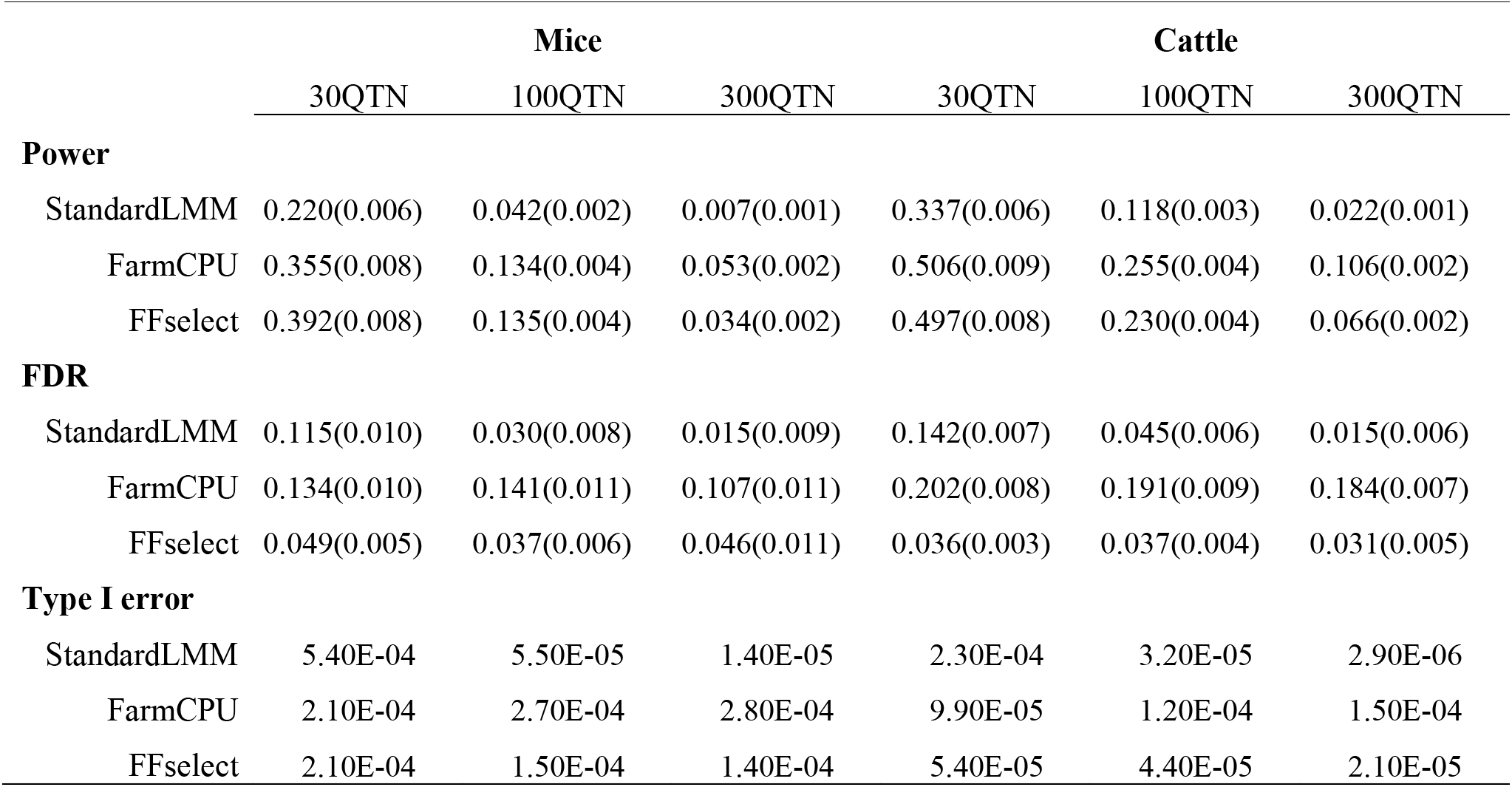
Comparison of statistical power, FDR, and type I error for mice and cattle genotype data with a simulated phenotype including environmental variance=0.1, genetic variance=0.5, and residual variance=0.4. Additive genetic effects were simulated with 30, 100, and 300 QTN. The statistical power, FDR, and type I error rate shown here are the averages of 100 simulation replicates (standard error of the mean indicated in parentheses).

|  | Mice |  |  | Cattle |  |  |
| --- | --- | --- | --- | --- | --- | --- |
|  | 30QTN | 100QTN | 300QTN | 30QTN | 100QTN | 300QTN |
| **Power** |  |  |  |  |  |  |
| StandardLMM | 0.220(0.006) | 0.042(0.002) | 0.007(0.001) | 0.337(0.006) | 0.118(0.003) | 0.022(0.001) |
| FarmCPU | 0.355(0.008) | 0.134(0.004) | 0.053(0.002) | 0.506(0.009) | 0.255(0.004) | 0.106(0.002) |
| FFselect | 0.392(0.008) | 0.135(0.004) | 0.034(0.002) | 0.497(0.008) | 0.230(0.004) | 0.066(0.002) |
| **FDR** |  |  |  |  |  |  |
| StandardLMM | 0.115(0.010) | 0.030(0.008) | 0.015(0.009) | 0.142(0.007) | 0.045(0.006) | 0.015(0.006) |
| FarmCPU | 0.134(0.010) | 0.141(0.011) | 0.107(0.011) | 0.202(0.008) | 0.191(0.009) | 0.184(0.007) |
| FFselect | 0.049(0.005) | 0.037(0.006) | 0.046(0.011) | 0.036(0.003) | 0.037(0.004) | 0.031(0.005) |
| **Type I error** |  |  |  |  |  |  |
| StandardLMM | 5.40E-04 | 5.50E-05 | 1.40E-05 | 2.30E-04 | 3.20E-05 | 2.90E-06 |
| FarmCPU | 2.10E-04 | 2.70E-04 | 2.80E-04 | 9.90E-05 | 1.20E-04 | 1.50E-04 |
| FFselect | 2.10E-04 | 1.50E-04 | 1.40E-04 | 5.40E-05 | 4.40E-05 | 2.10E-05 |

**Figure 1.**
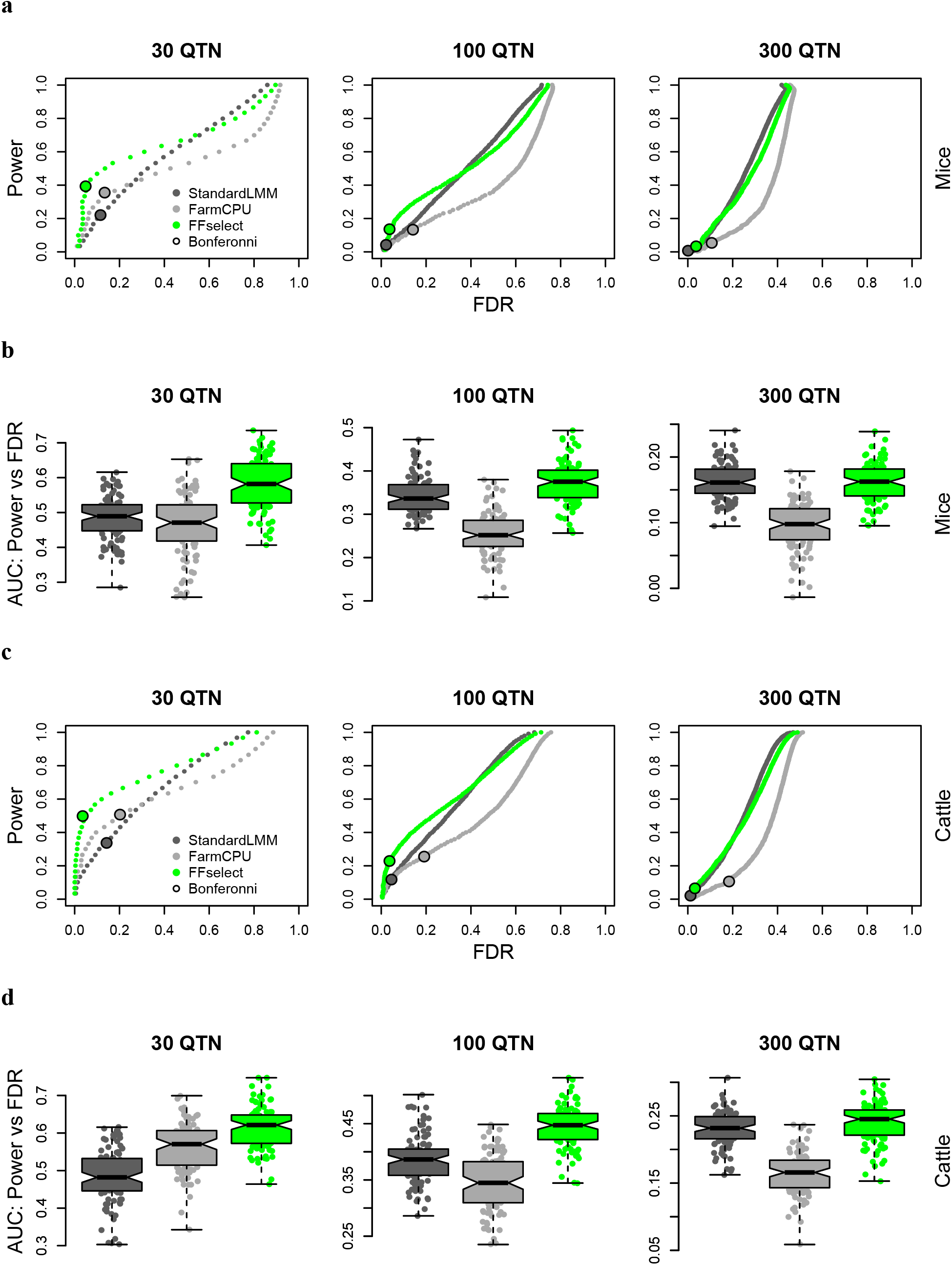
Mean statistical power plotted against false discovery rate (FDR) for a 30, 100, and 300 quantitative trait nucleotide (QTN) model for 100 simulation replicates. Three mapping methods were compared: Standard linear mixed model (standardLMM), FarmCPU, and FFselect in (**a, b**) mice, and (**c, d**) Holstein dairy cattle. A QTN was considered detected if a SNP within 2.5 MB on either side was declared significant. Boxplots of area under the curve (AUC) for power vs FDR for each of the 100 simulation replicates in (**b**) mice, and (**d**) Holstein dairy cattle.

FarmCPU and FFselect both demonstrate increased mapping precision (Figure 2) due to the fact that these methods take into account SNP(s) in LD with the causal SNP at a particular locus. FarmCPU models these SNP (pseudo QTN) as fixed effects in the final GWAS model whereas precision for the FFselect method is determined by a user specified window size for exclusion of pseudo QTN from the featured SNP relationship matrix (RM2).

**Figure 2.**
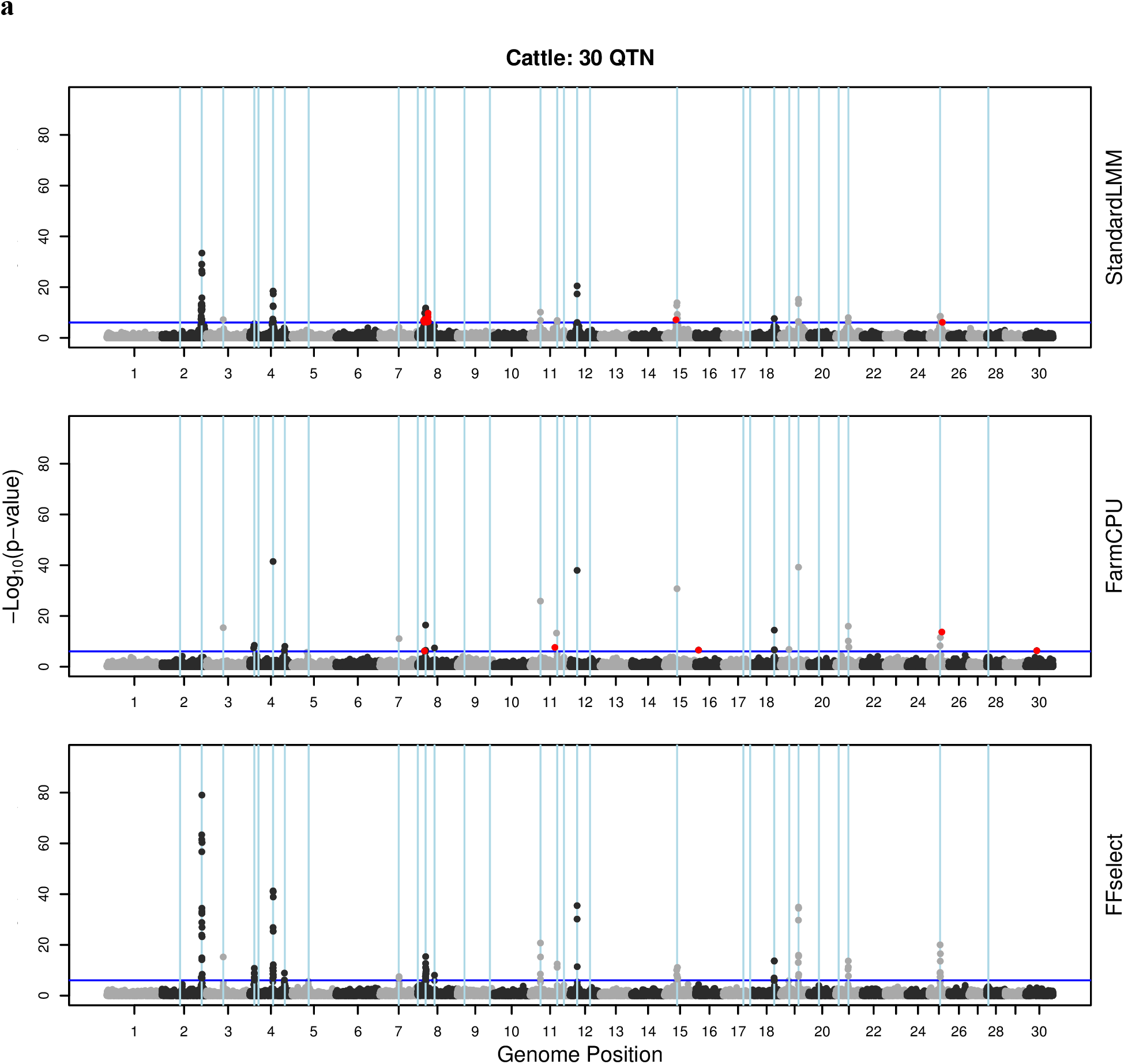
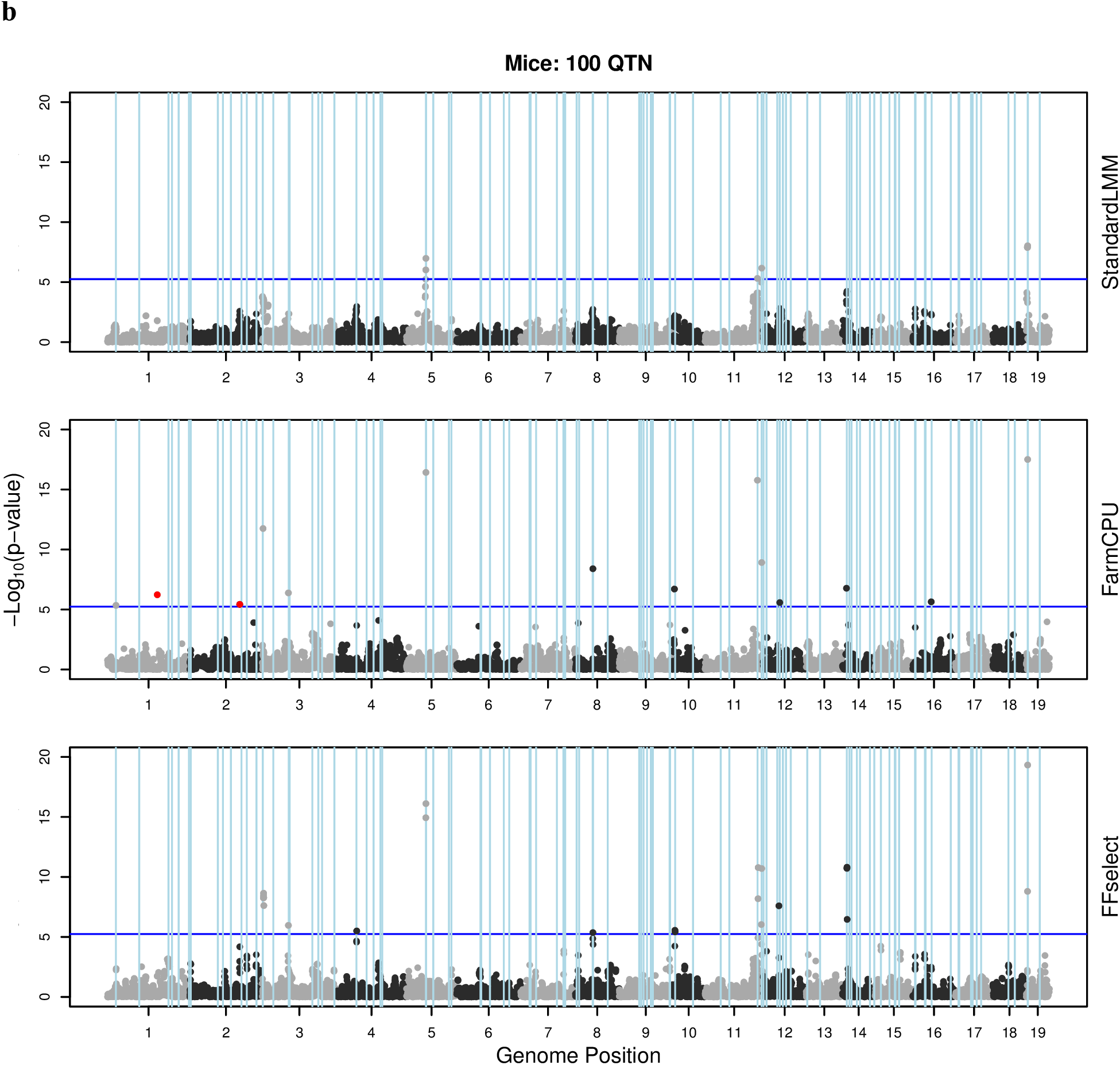
Representative examples of Manhattan plots for GWAS results for the standard linear mixed model (standardLMM), FarmCPU, and FFselect mapping methods selected form the 100 simulation replicates for a 30 quantitative trait nucleotide (QTN) model in (a) Holstein dairy cattle, and (b)100 QTN model in mice. Vertical lines indicate the position of true QTN. Red dots indicate false positives.

Genomic inflation of p-values for the different mapping methods were compared by examining quantile-quantile plot and genomic inflation factors (Figure 3). The FarmCPU method exhibited inflation of p-values in mice and cattle (with the exception of the 30 QTN model in mice) whereas the standardLMM and FFselect method genomic inflation factors did not deviate from a nominal false positive rate.

**Figure 3.**
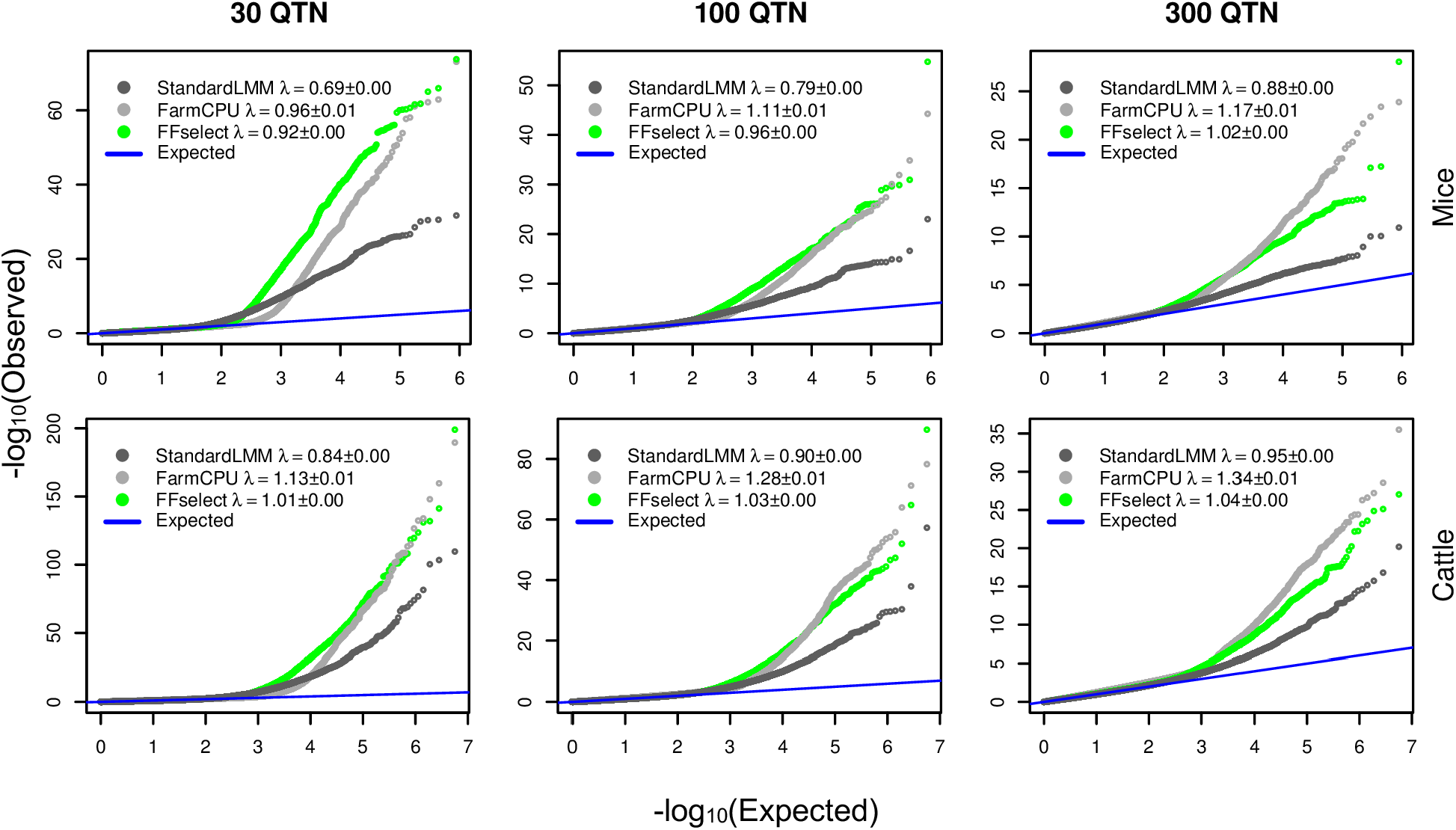
Observed versus expected distribution of p-values (QQ plots) for the standard linear mixed model (standardLMM), FarmCPU, and FFselect mapping methods in mice and Holstein dairy cattle. P values were from association tests on a simulated trait with environmental variance=0.1, genetic variance=0.5, and residual variance=0.4 controlled by 30, 100, or 300 quantitative trait nucleotides (QTN). QQ plots display results from 100 simulation replicates. λ=mean genomic inflation factor ± standard error of the mean for 100 replicates.

Computational time on a 2.3 GHz Intel Core i7 MacBook Pro for the 300 QTN model in the mice dataset comprised of 1814 individuals and 8850 SNP averaged 0.2, 0.6, and 6.2 minutes for the standardLMM, FarmCPU, and FFselect methods, respectively. Likewise, average computational time for the cattle dataset (3417 individuals, 56483 SNP) averaged 4.1, 3.7, and 9.2 minutes.

## Discussion

In addition to influence from the environment, complex traits are controlled by genetic variants of large and small effect. A statistical model that best captures the genetic architecture and environmental influences on a trait will improve mapping power while simultaneously controlling the false discovery rate. When testing for an association of test markers with the phenotype, FFselect includes an estimate of: 1) large effect genetic variants, 2) polygenic effects of small effect loci, and 3) shared environmental effects.

As demonstrated by simulations, FFselect displays improved performance in terms of power and FDR in comparison with methods that do not accurately model the effect of large effect loci (standardLMM) or small effect loci (FarmCPU). FarmCPU has previously been shown to greatly improve power over the standardLMM and other multi-locus GWAS approaches. In our simulations, FFselect and FarmCPU performed similarly in terms of power for traits controlled by a smaller number of loci and FarmCPU outperformed FFselect for traits controlled by a large number of loci, although at the cost of a high false discovery rate. FFselect demonstrated improved power over the standardLMM while adequately controlling the false discovery rate likely as a result of the inclusion of both a select featured SNP GRM in addition to a GRM comprised from all of the SNP. (Widmer et al., 2014) demonstrated the importance of including a mixture of two GRMs, one constructed from all SNP and another constructed from SNP identified by SNP selection. This approach yielded more power and better controlled type I error in the presence of population structure and familial relatedness. The stepLMM method described by (Li et al., 2017) includes a GRM comprised of all SNP and shares similarity with the FFselect approach; however stepLMM does not offer the option of including shared environmental effects.

FFselect allows for the modeling of shared environmental effects, a feature that is currently missing from most GWAS software and problematic in scenarios were genomic relationships are confounded by shared environment. FFselect partitions the phenotypic variance into large genetic effects, small genetic effects, and environmental effects, which allows the user to obtain estimates of the environmental variance, the explained and unexplained heritable variance. In addition, FFselect provides insight into trait genetic architecture based on the number of loci with larger genetic effects.

The mice dataset utilized in this study features full-sibs sharing the same cage, an extreme example of genomic relationships confounded by shared environment. Livestock datasets are often characterized by similar confounding, albeit to a lesser degree. For example, within dairy herds progeny are retained within the same herd and a limited number of sires are used for mating via artificial insemination resulting in increased genetic similarity within a herd confounded by shared environmental influences. Although only animal datasets were included in the current study, FFselect should also be extendable to human GWAS where household or geographic region may constitute a shared environment. Moreover, the FFselect environmental relationship matrix is not limited to being comprised of only zeros and ones with ones indicating a shared environment. The environmental relationship matrix could also be a distance matrix indicating spatial relationships.

Plotting of p-values by genome position for the FFselect method most closely resembles a “Manhattan” skyline in comparison to the other two methods we compared in this study. The standardLMM method resulted in poor mapping precision and SNP several megabase pairs distance from a causal marker of large effect were often reported to be significant. For this reason, we found it necessary to specify a 2.5 MB distance threshold from the causal marker when calling a hit a true positive. Without this relaxed distance threshold, the standard linear mixed model would have performed poorly in comparison to the FarmCPU and FFselect method. The FarmCPU plots, on the other hand, appear very much the opposite as a result of modeling all of the pseudo QTN as fixed effect covariates. The authors describe the FarmCPU plots as appearing more like “helicopters over Manhattan, Kansas” (Liu et al., 2016). The FFselect method allows users to specify the level of mapping precision by providing a threshold distance from a test marker in which pseudo QTN are excluded from the featured SNP relationship matrix. If the user specified a distance of 1 base pair the plots would look very similar to the FarmCPU plots whereas specifying a distance of 5 MB would produce plots more similar to the standardLMM method.

A criticism of feature selection methods is the potential to overfit the data, especially when selection of features is performed and applied within the same dataset. Ideally, it would be preferable to have a separate training set for selection of pseudo QTN and a test dataset to optimize the weight (contribution to the explained variance) of the featured SNP relationship matrix and to perform the GWAS. This approach, however, would require a larger sample population size. In an effort to prevent overfitting using the FFselect method, we included an additional criteria (BIC) to stop the forward feature selection of pseudo QTNs. Alternatively, implementation of Bayesian approaches for the selection of pseudo QTN could be examined, although this type approach may be computationally less feasible for large datasets.

In summary, FFselect has built upon recent advances that improved the power of the linear mixed model for genome-wide association analysis and extended these methods to control for environmental confounding in a computationally feasible approach. Computational time for the FFselect method was slightly longer in comparison to other methods used in this study; however it is worth noting that the FFselect R function relies heavily upon the lrgpr R package (Hoffman et al., 2014) which allows analysis of large datasets in parallel on multicore computers and which is designed to take advantage of the bigmemory R package (Kane, Emerson, & Weston, 2013) for out-of-core computing to efficiently process datasets too large to fit into main memory. Additionally, the lrgpr R package is able to perform GWAS models that include SNP interaction terms. The call to lrgprApply when performing the GWAS within the FFselect function can be easily modified to allow test marker effects to vary by environment (provided sufficient number of individuals per environment) or any other user specified covariate (e.g. y ~ SNP + SNP:environment, instead of y ~ SNP).

## Acknowledgements

Parts of this research first appeared in the thesis of NS (Schultz, 2016).

## Author Contributions

NS conceived the study, performed the analysis, and drafted the manuscript. KW contributed to editing of the manuscript. All authors approved the final manuscript.

## Data Availability Statement

R code for the FFselect method and a provided example utilizing the mouse dataset are available at https://github.com/NicholSchultz/FFSelect

The cattle data has been made publicly available in the form of data sharing with scientists from the USDA–Agricultural Research Service and Council on Dairy Cattle Breeding (Bowie, MD)

